# Inter-subject pattern analysis for multivariate group analysis of functional neuroimaging. A unifying formalization

**DOI:** 10.1101/2020.08.28.272153

**Authors:** Qi Wang, Thierry Artières, Sylvain Takerkart

## Abstract

**Background and objective:** In medical imaging, population studies have to overcome the differences that exist between individuals to identify invariant image features that can be used for diagnosis purposes. In functional neuroimaging, an appealing solution to identify neural coding principles that hold at the population level is inter-subject pattern analysis, i.e. to learn a predictive model on data from multiple subjects and evaluate its generalization performance on new subjects. Although it has gained popularity in recent years, its widespread adoption is still hampered by the blatant lack of a formal definition in the literature. In this paper, we precisely introduce the first principled formalization of inter-subject pattern analysis targeted at multivariate group analysis of functional neuroimaging.

**Methods:** We propose to frame inter-subject pattern analysis as a multi-source transductive transfer question, thus grounding it within several well defined machine learning settings and broadening the spectrum of usable algorithms. We describe two sets of inter-subject brain decoding experiments that use several open datasets: a magnetoencephalography study with 16 subjects and a functional magnetic resonance imaging paradigm with 100 subjects. We assess the relevance of our framework by performing model comparisons, where one brain decoding model exploits our formalization while others do not.

**Results:** The first set of experiments demonstrates the superiority of a brain decoder that uses subject-by-subject standardization compared to state of the art models that use other standardization schemes, making the case for the interest of the transductive and the multi-source components of our formalization The second set of experiments quantitatively shows that, even after such transformation, it is more difficult for a brain decoder to generalize to new participants rather than to new data from participants available in the training phase, thus highlighting the transfer gap that needs to be overcome.

**Conclusion:** This paper describes the first formalization of inter-subject pattern analysis as a multi-source transductive transfer learning problem. We demonstrate the added value of this formalization using proof-of-concept experiments on several complementary functional neuroimaging datasets. This work should contribute to popularize inter-subject pattern analysis for functional neuroimaging population studies and pave the road for future methodological innovations.

## 1. Introduction

One of the major challenges encountered in medical signal and image processing is to overcome the large heterogeneity often present in the data. This heterogeneity can be introduced by differences that exist across acquisition devices, across modalities, across populations or across individuals (see e.g. [6], [3]). In functional neuroimaging group analysis, the objective is to unravel neural coding principles that are invariant throughout a given population, or that differ across groups of individuals, e.g. between patients and healthy subjects. Since the 2000s, the advent of multivariate pattern analysis has opened new opportunities for this at the macroscopic level, by rendering explicitly usable the information that lies in the differential modulations of brain activation across multiple locations – i.e. multiple sensors for electro-encephalography (EEG) and magneto-encephalography (MEG), or multiple voxels for functional magnetic resonance imaging (fMRI) – through the use of machine learning algorithms (see reviews in e.g. [27, 19]). A straightforward scheme to examine these modulations throughout a population is to perform inter-subject pattern analysis (ISPA), i.e. to train a predictive model on data recorded in a set of subjects and test it on data from new subjects [44]. Indeed, obtaining an above-chance generalization performance in such context indicates that the algorithm has implicitly identified neural coding principles that hold across all individuals of the population.

Despite the fact that ISPA offers a direct way to assess neural coding principles at the group level, the most commonly used method for population studies of multivariate brain activation patterns is an alternative hierarchical strategy based on learning individual models, each from data recorded in a single subject, and aggregating their results in a second level analysis (see e.g. [37]). We believe that this is the case for two main reasons. First, as reported early on, e.g in [20, 12], and more recently discussed, e.g in [19], it is more difficult to obtain good generalization performances on data from new subjects because of the existence of interindividual variability, which can make within-subject multivariate pattern analysis more immediately rewarding for the neuroscientist. Furthermore, having access to subject-level results can also facilitate the interpretation in this case. Secondly, a formal definition of ISPA is lacking, which is an obstacle to design dedicated algorithms that could be implemented in standardized methods and offered to the community in user-friendly software packages.

Having recently demonstrated the potential advantages of ISPA over the standard hierarchical approach (see [44]), we here compensate for the latter by introducing a principled formalization of ISPA. First, we precisely define the ISPA setting as a solution for group-level multivariate analysis of functional neuroimaging data, and we frame it as a *multi-source transductive transfer* learning problem, thus using a combination of well defined machine learning concepts (Section 2). Then we detail two experimental studies that demonstrate the added value of this formalization (Section 3). Finally, we describe how it can offer a unifying framework for existing ISPA algorithms and we discuss several potential avenues to address the remaining methodological challenges raised by ISPA (Section 4).

## 2. Material and methods

### 2.1. Machine learning reminders

First, we provide the definitions of several machine learning concepts that we will exploit to formalize Inter-Subject Pattern Analysis as a multivariate group analysis method for functional neuroimaging experiments.

#### Transfer learning

One of the main hypotheses used in standard machine learning is that all the data live in the same feature space and follow a single probability distribution. In particular, in order to ensure that a model *f* estimated on some training data will be able to generalize to other data (i.e a test data set), one needs that the feature spaces and distributions of the training and test data sets are the same. However, in numerous real world applications, this is not the case, often because one wishes to use the model in a slightly different context than the one within which it was estimated, i.e because of what is called a *dataset shift* between the training and testing phases. In such case we hope that the information learnt in one context can be *transferred* to another one, which defines the *transfer learning* problem [31]. Furthermore, according to the nomenclature described in [39] to characterize the different dataset shifts that can occur in a transfer learning setting, one refers to a *domain shift* when the feature spaces are different and to a *distribution shift* when the probability densities are different.

#### Transduction and transductive transfer

The term *transduction*, introduced in [16] as an alternative to the more classical inductive setting, designates a particular case of supervised machine learning where several – or all – the data points of the test set are available when the model is trained, without their labels. Several usages of transductive inference can be encountered. First, when one only wishess to obtain labels for the samples of the test set without the need for a predictive model that can further generalize to future new data, one can exploit the full test set to guess these labels, for instance by performing label propagation from the training to the test set [48]. Secondly, when dealing with a transfer learning problem, exploiting a set of unlabeled samples of the test set can allow improving the generalization capability of a transfer model. Methods that exploit this opportunity are tagged as *transductive transfer* methods since [1], a nomenclature that has been taken up in the survey presented in [31] and widely adopted since.

#### Multi-source learning

In most transfer learning problems, the model is trained on data obtained in a single context, and the main challenge is to transfer the knowledge to a different one for the test data. However, it is also possible that the training set itself gathers data obtained in multiple contexts instead of a single one. One then wishes to take the context information into account during the training of the model, a challenge that has been formalized as *multi-source learning* [13]. Note that the term *multi-source* appears in the literature with very different meanings, sometimes designating heterogeneous data, or being used as a synonymous for multiview or multi-modal data etc. We insist on the fact that we here use the clearly defined multi-source learning setting proposed in [13], which unambigusously designates the supervised learning problem where a single model has to be learnt from training data drawn from several probability distributions but associated with a common output space 𝒴.

### 2.2. A generative model of functional neuroimaging data

We assume that we have at hand the data recorded during a functional neuroimaging experiment that was performed by a group of individuals drawn from a single population, who each performed a set of tasks over numerous trials. Let 𝒮 = {1, …, *S*} be the set of subjects who participated in this experiment. For subject *s*, let 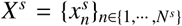 be the dataset, where *N*^*s*^ >> 1 is the number of samples available for subject *s* (which can be the number of timepoints of the time-series, or more classically the number of trials performed by the subject). Each of these samples 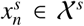 is associated with a value 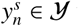 which characterizes what was done by the subject while 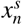 was recorded. The *y* variable can be discrete (e.g. face vs. house stimuli) or continuous (e.g. the reaction time), thus defining a classification or a regression problem respectively. The full dataset *D* is given by: 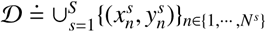. A generative model of this data consists in stating that for a given subject *s*, the pairs 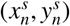 are realizations of a probability distribution 𝒫^*s*^. Following these definitions, the critical points to be noted are that i) 𝒴 is common to all subjects because they participated in the same experiment and performed the same tasks, ii) 𝒳^*s*^ can differ across subjects since the brain of each individual is unique (for instance, the size of the primary visual cortex can vary by a factor greater than two across healthy individuals [38]), iii) 𝒫^*s*^ can also depend on *s* because the noise level in the data and the properties of the signal (e.g. the amplitude and location of informative features) often varies across subjects. All these properties of the dataset 𝒟 are illustrated on Fig. 1.

**Figure 1:**
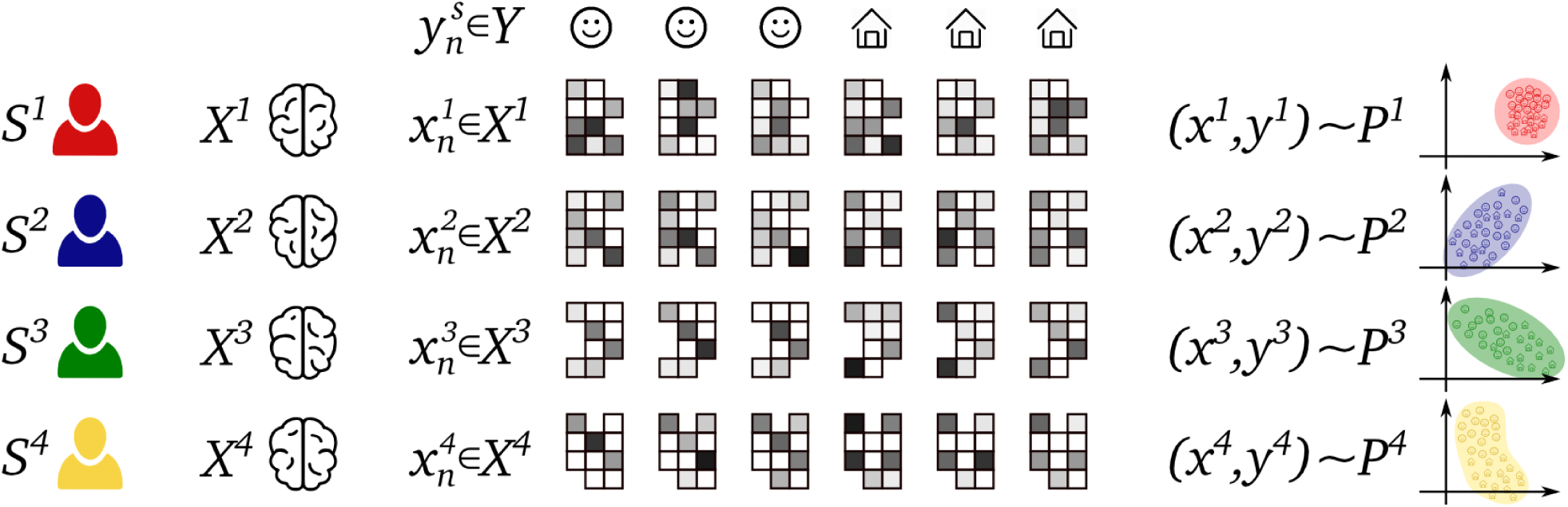
Illustration of the data at hand when performing inter-subject pattern analysis and the associated challenges. Four subjects (*S* ^1^, *S* ^2^, *S* ^3^, *S* ^4^) are depicted, each with their respective data domain 𝒳^*s*^ defined by their individual brain. They have partcipated in the same experiment (for instance looking successively at a face or a house during each trial), which defines the output variable 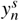 in a space that is common to all participants, while their brain activity 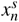 is recorded (here, the different rectangle-like grids represent brain patterns recorded in a brain area that is homologuous across subjects, yet its shape and size vary across subjects, as with, e.g., the primary visual cortex [38]). The data from each subject follows its own probability distribution 𝒫^*s*^, as illustrated in a putative two-dimensional feature space on the right.

### 2.3. Defining inter-subject pattern analysis for group multivariate analysis

We now define precisely what we denote as Inter-Subject Pattern Analysis (ISPA) when used for multi-variate group analysis of functional neuroimaging data. ISPA is a group analysis framework that aims at assessing neural coding principles associated with the tasks performed by the subjects at the population level, at the spatial resolution offered by the imaging modality available. It consists in designing a model that can predict a trait associated with the experimental paradigm – such as the category of a stimulus or a reaction time – from a multivariate pattern of brain activity. For this, a machine learning model is trained on data from a subset of subjects 𝒮_*train*_ and its generalization power is measured on the left-out subjects 𝒮_*test*_. In order to provide a robust assesment of neural coding principles over the full population, including for subjects not present in 𝒮, this operation is repeated with different splits of train- and test-subjects using cross-validation schemes that follow a leave-*P*-subjects-out rule – the most common one being the leave-one-subject-out scheme. Note that if *P* > 1 and if the number of subjects *S* is large, this yields a very large number of splits; in such case, one can randomly select a subset of splits to reduce the computational burden [43]. The most typical usecase of ISPA is inter-subject brain decoding, where a classifier is trained on data from a set of subjects and evaluated on others in order to identify features that are consistently involved in the cognitive processes implicated in the tasks performed by all the individuals of the population (see e.g. [21]).

In order to better define ISPA and narrow down its specificities, we now describe its commonalities and differences with two closely related problems encountered in medical signal and image analysis.

#### Link with Computer-Aided Diagnosis systems (CAD)

As in ISPA, CAD systems aim at performing predictions on data from new individuals [40]. However in CAD, the prediction targets a variable that directly characterizes the patient and is performed from a single observation, as e.g. when a diagnostic status is guessed from the shape of the patient’s brain measured with anatomical MRI, or when the severity of the disease is inferred from the patient’s functional connectome. In ISPA, the output variable does not characterize the subject, but each experimental trial performed by the subject, and many predictions are produced on the data from each individual.

#### Link with Brain-Computer Interface (BCI)

When designing a BCI [9], the underlying model can be trained on calibration data from the subject who will use the interface and/or on data from several individuals, making BCI design analogous to ISPA in this latter case. However, while in BCI, it is the performance of the device on any single trial that matters, in ISPA, the population-wise consistency of the neural coding principles is assessed from the predictions on *all* the trials of a test subject. A direct corollary is that in BCIs, the model needs to be accurate also on future events and can be progressively updated in an *online* manner, whereas in ISPA the entire dataset is available at the time of the analysis and it will not grow in the future: it is a purely *offline* learning problem.

### 2.4. ISPA: a multi-source transductive transfer problem

For a given split of 𝒮 into 𝒮_*train*_ and 𝒮_*test*_, ISPA implies training a model on data from 𝒮_*train*_ and assessing its generalization performance on subjects in 𝒮_*test*_. Obtaining predictions that are close to the real targets 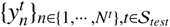 would provide evidence that the modulations of brain activity with respect to the values of *y –*i.e. the coding of *y* in brain patterns *X* – are consistent across training and test subjects, and hence throughout the full population from which these subjects were drawn. Given the previously defined concepts and definitions, we put forward the following claims:

1. first, because the test set is composed of data from new individuals, thus drawn from distribution(s) and recorded from brain(s) (i.e feature spaces) not represented in the training set, ISPA raises a *transfer learning* question; furthermore, ISPA combines the challenges raised by both domain shifts and distribution shifts;
2. secondly, because group analyses are performed after the recording of the data for all the subjects that participate in a study (i.e in an offline setting), and because inference on brain function at the group level can only be drawn from the prediction of *all* labels of the test subject(s), ISPA can be considered in a *transductive* context, i.e it is valid to exploit all data from the test subject(s), without their labels, to train the model;
3. lastly, the nature of the training set, which gathers data from different participants, drawn from their own distribution 𝒫^*s*^ and available in their own feature space (i.e their brain 𝒳^*s*^), makes of ISPA a *multi-source learning* problem, i.e. each participant provides a *source* of data.

Accordingly, ISPA can be formalized as a *multi-source transductive transfer* learning problem, thus combining three well defined machine learning concepts. Although the combination of two of these three concepts is fairly common in the literature (see e.g. transductive transfer [31], or multi-source domain adaptation [25]), designing methods that exploit the full extent of these three concepts remains an important challenge in current machine learning research (as met, e.g., in natural language processing when different languages are modelled as providing data from multiple sources). The rest of the paper aims at validating the relevance of this formalization using an experimental approach, as well as discussing the opportunities that this formalization opens for future methodological developments.

## 3. Experiments and results

In this section, we describe two sets of original experiments. The objective of these experiments is to assess the relevance of each of the three components of our multi-source transductive transfer setting for examining the challenges raised by ISPA. Importantly, note that we do not aim here at designing a new machine learning algorithm that will beat existing ones. We therefore exploit algorithms which are commonly used by neuroscientists and offer performances close to the state-of-the-art in brain decoding experiments.

### 3.1. Experimental data

We use functional neuroimaging data from two previously published studies. The first one includes 100 participants from the fMRI data used in [32] (data available at http://openneuro.org/datasets/ds000158/) to identify voice sensitive areas in the temporal cortex, using a *voice localizer* paradigm [5]. The participants passively listened to 40 blocks of auditory stimuli (20 vocal blocks and 20 non-vocal blocks). The data processing pipeline included the co-registration of the functional data with the T1 anatomical image, the correction of motion in the fMRI images and the estimation of a general linear model that included one regressor for each of these blocks with additional regressors of non intestest such as motions parameters. The corresponding regression coefficients (beta maps) provide estimates of the single-block functional response. These operations were performed using SPM12 [2]. Then, using the *freesurfer* software suite [14], the T1 image was processed to estimate the three-dimensional cortical mesh, onto which each beta map was projected and then resampled onto the *fsaverage* template. An anatomical region of interest (ROI) was defined to englobe the auditory cortex as well as voice sensitive regions, separately in each hemisphere, using the Desikan parcellation provided in freesurfer. Since standard multivariate analysis are performed in a single contiguous brain area, we defined two sets of data, hereafter denominated “fMRI 1” for the left hemisphere and “fMRI 2” for the right hemisphere, as is often done to study lateralization effects. The task was to decode whether the participant had heard vocal or non-vocal stimuli from the brain responses recorded in each of these two regions. The second study is the event-related MEG experiment from [22] (data available at https://www.kaggle.com/c/decoding-the-human-brain), where 16 participants viewed one picture per trial, either of a face or of a scrambled face. On average, 580 trials were available per subject. In order to construct our input feature vector, the raw data was processed using the MNE-Python software [17]: we extracted the timeseries from a 500*ms* window after stimulus onset and we temporally downsampled the data by a factor of eight. This provided us with a third dataset, hereafter simply denominated “MEG” data. The task of the decoder was to guess whether the subject was viewing a real face or a scrambled one from the MEG data. Interestingly, these datasets present complementary characteristics – their nature (fMRI vs. MEG), the studied domain (within ROIs vs. on the full brain), the nature of the paradigm (block vs. event-related), the number of subjects available (100 vs. 16), the number of samples available for each subject (40 vs. around 580).

### 3.2. General experimental setting

In all experiments, we use cross-validation schemes where several subjects are left out for the test set in each data split (leave-*P*-subjects-out) – the exact scheme being detailed hereafter in each case. The generalization performance of the model is assessed with the average classification accuracy across all splits, which offers a robust and unbiased estimate thanks to the large number of splits available [43]. We chose to use two families of classifiers, logistic regression and linear support vector machine (SVM) for the following reasons: i) they are simple to use and easily available, ii) they have become the de facto standards in functional neuroimaging, and iii) despite a large body of work throughout the scientific community in the last decade, they still perform very close to the state of the art in brain decoding tasks [23] while remaining the most computationally efficient. The values of the hyper-parameters of the models are chosen through an inner cross-validation performed within the training set. For both types of models, SVM and logistic regression, the regularization weight *C* was chosen within {*10, 1, 0.1, 0.01, 0.005, 0.001, 0.0008, 0.0005, 0.0001, 0.00005, 0.00001*}. For logistic regression, we also selected the type of regularization (*l*_1_ or *l*_2_).

### 3.3. Experiment 1: assessing the interest of the multisource and transductive settings through feature standardization

*Aim*. In machine learning, the commonly used *feature standardization* operation consists in removing the mean – across samples – of each feature and scaling it to unit variance before feeding the data into the learning algorithm. We here aim at assessing the relevance of our formalization using this operation, by introducing standardization strategies that are multi-source (i.e aware of the multi-subject nature of the data) or not (i.e that simply pool the data from all subjects together), and others that are transductive or inductive. We therefore propose to benchmark four standardization strategies:

- a pooled inductive standardization: this is the classical machine learning strategy, the parameters are estimated on the full training set (which means, for ISPA, after a pooling of the data from all subjects in 𝒮_*train*_), and used to transform both the training set and the test set, item by item;
- a multi-source inductive standardization: in this case, for all the data of the training set, the standardization is performed independently on each subject; we then select the median training subject according to its standardization parameters, and transform all the test data points using the parameters learnt on the training set, item by item, i.e in an inductive manner;
- a pooled transductive standardization: as in the classical standardization (pooled inductive), all the training subjects are pooled together and standardized; but the test data is standardized independently on each subject, therefore exploiting all the unlabeled samples of each test subject: this is a transductive strategy;
- a multi-source transductive standardization: in this one, the standardization is performed subject by subject, whether in 𝒮_*train*_ or 𝒮_*test*_; this one is used in the literature, and made valid by our formalization.

These four strategies are formalized in greater details in Appendix A. This yields a 2 × 2 factorial design that will allow to independently evaluate the influence of the transduction and of the multi-source setting. For the sake of exhaustivity, we also included the case where no standardization is performed.

#### Experimental setting and results

With the fMRI data, we randomly defined 50 splits of the 100 subjects, each time including ten subjects for the test set and the other 90 for training. For the MEG data, we used a leave-two-subjects-out cross-validation scheme, which, for the 16 available subjects, yielded 120 train-test splits, which we all included in our analyses. The results of this first experiment are summarized on Fig.2. For each of the six cases (three datasets, two families of classifiers), we used a two-way analysis of variance to analyze the mean generalization performances of the estimated predictive models over all splits, across the four standardization strategies. It first revealed that the main effect of the *inductive vs. transductive* factor was very significant for all three datasets and for both logistic regression and SVM (*F* > 50.7 across all six cases, with *p* < 10^−11^), with greater performances for the transductive strategies in all cases. This demonstrates the added-value of the transductive setting for ISPA. Secondly, the main effect of the *pooled vs. multi-source* factor was not significant across all strategies. The interaction between the two factors was either significant or showed a trend towards significance (for the MEG and fMRI 2 datasets: *F* > 3.57, *p* < 0.05; for the fMRI 1 dataset: *F* = 2.51, *p* = 0.11 for logistic regression, *F* = 2.45, *p* = 0.12 for SVM). When targetting the better-performing transductive setting, a post-hoc paired *t*-test between the pooled and the multi-source transductive standardizations demonstrated the superiority of the multi-source transductive standardization (*t* > 3.49 across all six cases, *p* < 2.10^−3^). Finally, as a sanity check, we tested whether the models trained from data that was pre-processed with multi-source transductive standardization offered higher generalization performances than the models trained with the raw un-standardized data; the result was highly significant in all cases (paired *t*-tests, *t* > 12.6, *p* < 10^−16^).

**Figure 2:**
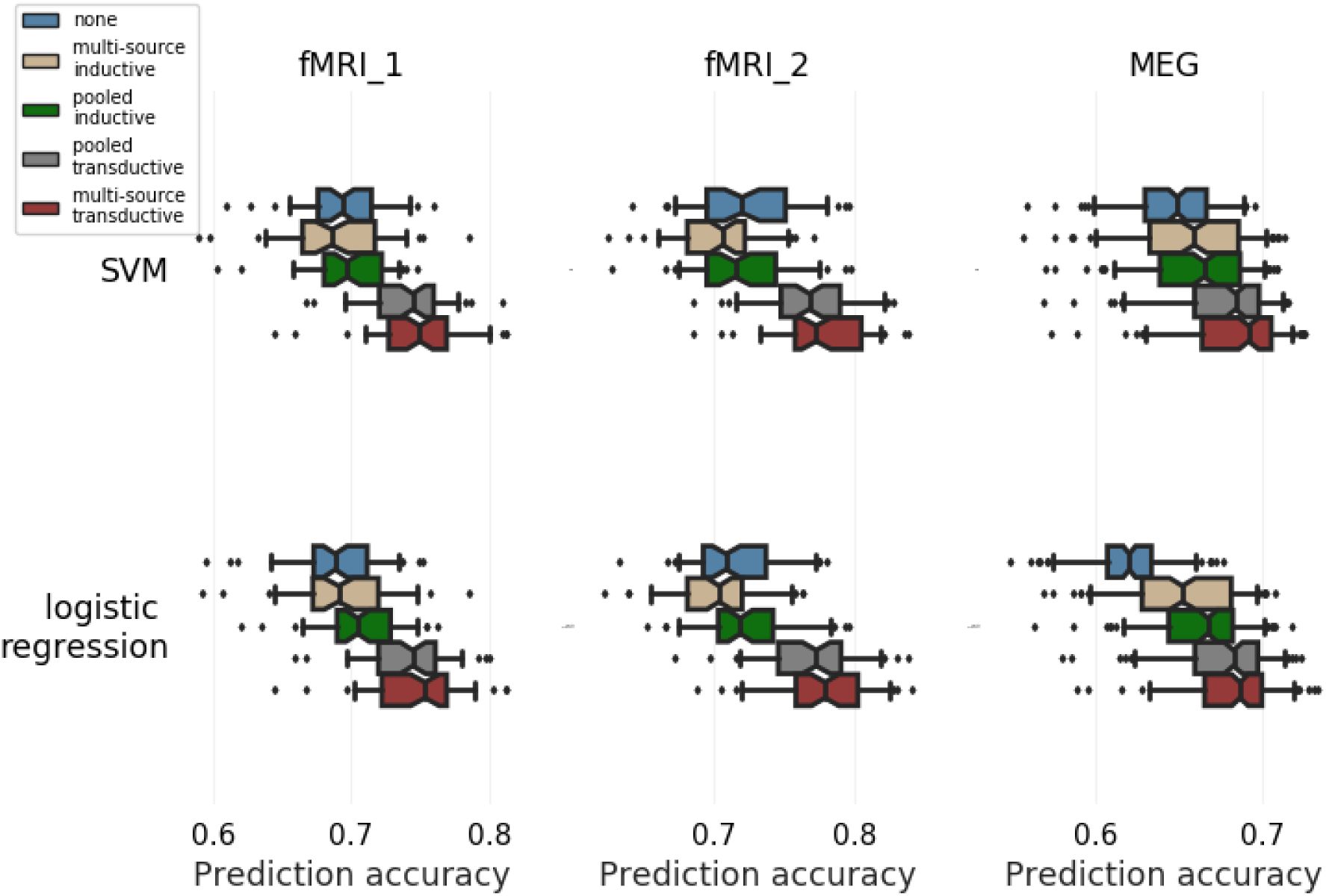
Experiment 1: Studying the effect of the transductive and/or multi-source settings on the quality of the inter-subject predictions, using feature standardization. The chance level for prediction accuracy is 0.5. First, the transductive strategies significantly outperform the inductive ones (see text). Secondly, in the transductive setting, the multi-source transductive standardization performs significantly better than the pooled transductive setting. Overall, the subject-by-subject multi-source transductive standardization offers the best prediction performances, using both logistic regression and SVM.

#### Conclusion

In the context of ISPA, our formalization validates the practice that consists in performing feature standardization on a subject-by-subject basis, which corresponds to the multi-source transductive strategy benchmarked here. The results described in the present experiment demonstrate that i) transduction, i.e having access to all the data points of the test subject(s), allows boosting the generalization performance of the model, ii) the multi-source setting provides an additional gain in performance when combined with transduction, and iii) the subject-by-subject standardization offers the best performances of all the data pre-processing strategies assessed here.

### 3.4. Experiment 2: measuring the transfer gap

#### Aim

Because it is likely to observe strong dataset shifts across individuals (as evidenced in e.g [20, 12]), obtaining a model that is valid for the whole population is a transfer learning question. We here perform original experiments that make it possible to quantify the transfer gap across subjects. Importantly, these experiments are conducted after multi-source transductive standardization, which allows for a first attenuation of the between-subject shifts. We reason as follows: if the between-subject dataset shifts were neglectable, the data of all subjects would follow a single probability distribution, and a decoder would perform identically on any new data, whether from subjects in 𝒮_*train*_ or from new subjects unseen at training time. *Experiment 2* precisely aims at probing this hypothesis by training a single model and evaluating its performances on two sets of new data, taken either in new subjects or from independent samples of the training subjects.

#### Experimental setting

First we extract two disjoint subsets of subjects 𝒮_1_ and 𝒮_2_ (of sizes *ψ*^1^ and *ψ*^2^ respectively) from 𝒮. For each subject *s* in 𝒮_1_, we split the data into 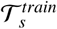 and 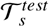. We use 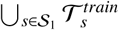 as the training set. We then define two different test sets: 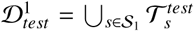, composed of unseen samples from training subjects, and 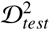 which gathers the data from the new subjects of 𝒮_2_. Moreover, we repeat this with different values of *ψ*^1^ to study the effect of the size of the training set. In practice, since the fMRI datasets include a large number of subjects we first define 𝒮_2_ by selecting a fixed set of *ψ*^2^ = 50 subjects. We then define 𝒮_1_ amongst the 50 other subjects, by randomly selecting *ψ*^1^ subjects (with *ψ*^1^ ∈ {10, 20, 30, 40}), and repeat this 30 times with different random draws. For the MEG dataset, the combinatorial possibilites are much more limited because it includes only 16 subjects. We work with a set of *ψ*^2^ = 2 subjects in 𝒮_2_, which we will repeatedly draw randomly within the 16 subjects. Then, for each of these draws, we define 𝒮 _1_ within the leftover subjects, with sizes *ψ*^1^ ∈ {5, 7, 9, 11, 13}. For each value of *ψ*^1^, we therefore obtain 50 measurements of the model accuracy. In all cases, we perform subject-by-subject feature standardization (as in *Experiment 1*) before fitting the model.

#### Results

The results, shown on Fig.3, are examined using an analysis of variance (ANOVA) with two factors –the nature of the test set (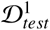 or 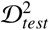) and the number of subjects *ψ*^1^ in the training set. The main effect of the number of subjects was found to be significant in all three cases (*F* > 20.96, *p* < 10^−11^), showing that the predictive power of the model improves when data from more subjects are added to the training set, which was expected since it corresponds to an increased size of the training set [12]. More importantly, the main effect of the nature of the test set was also significant in all cases (*F* = 9.62, *p* < 10^−2^ for fMRI 1, *F* = 295.02, *p* < 10^−42^ for fMRI 2 and *F* = 1823.35, *p* < 10^−166^ for MEG), showing that it is indeed more difficult for the model to generalize to data from new subjects than to new data from the training subjects. This demonstrates that the between-subject dataset shifts cannot be ignored, even after multi-source transductive standardization. Beyond these very significant main effects, other more complex ones appear. First, we observe a decrease of the accuracy on 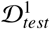 for the MEG data (green curve), which could be explained by the fact that the increased heterogeneity takes over the increased training set size, as reported and discussed in [35]. Secondly, the models offered different generalization performances between the left and right auditory cortices (fMRI 1 et fMRI 2 datasets respectively), which might be consistent with a recently reported asymmetry of the amount of inter-individual functional variability in the auditory cortex [34]. Both these results will deserve further investigations in the future.

**Figure 3:**
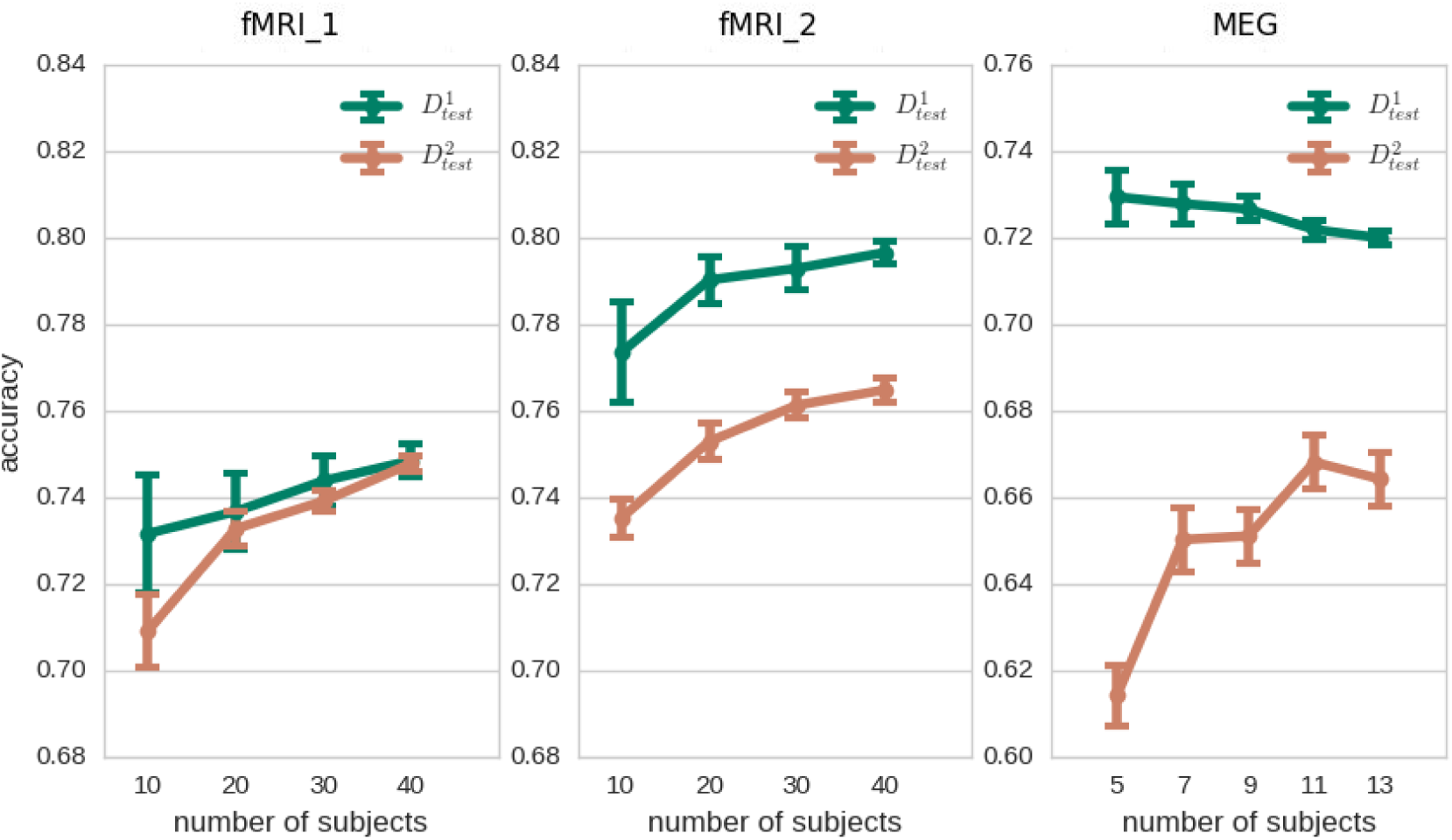
Experiment 2: Assessing the transfer gap. The accuracy of a given model (chance level is 0.5) is compared on two test sets composed of new data from the training subjects (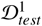, green curves) or data from new subjects (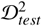, orange curves). Two effects are clearly visible: 1. increasing the sample size available for training by adding subjects in the training set allows improving the generalization to new subjects’ data (orange curves); 2. it is more difficult to generalize to data from new subjects. The latter demonstrates that we cannot ignore the distribution shifts between subjects, i.e. that pooling data across subjects while ignoring their differences is a sub-optimal strategy, even after multi-source transductive feature standardization.

#### Conclusion

In classical machine learning, one of the main assumptions is that the training and test sets contain data drawn from the same probability distribution. This experiment unequivocally demonstrates that this assumption is not met in ISPA experiments, even after the inter-subject dataset shifts have been attenuated using multi-source transductive feature standardization. It is therefore critical to thouroughly consider ISPA as a transfer learning question, but also to go beyond standard cross-subjects data pooling by taking into account the multi-subject nature of the training set, as put forward by our formalization.

## 4. Discussion

In this paper, we have proposed a definition of intersubject pattern analysis (ISPA) as a group analysis framework for multivariate pattern analysis of functional neuroimaging data, and formalized it as a multisource transductive transfer learning question. Furthermore, we have experimentally demonstrated the relevance of each of the three machine learning concepts that contribute to this formalization, i.e. the multisource and transductive settings through *Experiment 1*, and the transfer setting through *Experiment 2*. For the neuroscientist, the first direct benefit of this formalization is to theoretically warrant the use of any type of data transformation performed on a subject-by-subject basis, including on the test subjects, as long as the labels are not exploited. As a proof of concept, *Experiment 1* has shown that subject-by-subject standardization outperforms all others standardization strategies. We believe that it is the case because the effect of such subject-by-subject operation goes far beyond classical feature standardization, by attenuating the distribution shifts that exist between individuals: this yields both a more homogeneous training set – easing the learning task itself, and a test set that is closer to the training data – thus facilitating the generalization to new subjects. In fact, it is already common practice for ISPA studies. Indeed, it has been reported in several articles (see e.g. [29]; [36]; [24]; [11]; [8]; [46]), but we believe it is also vastly used without being reported because researchers might consider it to be trivial practice. Our formalization highlights that it is in fact not trivial, and we therefore recommend researchers to explicitly report in their publications whether they perform subject-by-subject feature standardization using this multi-source transductive setting.

We now attempt to put our formalization in perspective with respect to the existing literature of ISPA-dedicated methods. A full review of this literature is clearly out of the scope of the present paper, but we aim at providing illustrative examples of the different questions to be handled when faced with a multi-source transductive transfer learning question. The first challenge raised by ISPA is to handle the multi-source nature of the data, i.e. the fact that the data points hailing from multiple subjects originally live in different spaces 𝒳*s* (domain shift) and are drawn from different probability distributions 𝒫^*s*^ (distribution shift), as defined in Section 2 and illustrated on Fig. 1. The most basic solution for this uses image registration based on brain anatomy to bring the patterns *X*^*s*^ of each individual into a common space 𝒳 defined by a brain template such as the MNI. Although this practice is vastly used and has proved potentially effective for searchlight decoding [44], it is well known that it cannot fully overcome functional differences between individuals [42]. Numerous more advanced methods have since been proposed in the literature to overcome the between-subjects domain shifts by constructing a function-based template space X into which the data of all subjects is transformed: non-diffeomorphic function-based alignment methods [18], multiway canonical correlation analysis [10], projections into spatially-structured graph spaces [41] or riemanian spaces [4] etc. Their effectiveness is usually demonstrated by an improved inter-subject decoding performance. However, it has never been explicitly quantified to what extent these deterministic transformations allow reducing the between-subject shifts that exist between distributions {𝒫^1^, …, 𝒫^*S*^}; this clearly should be tackled to assess whether these approaches fullly resolve the challenge raised by between-subject dataset shifts. Other methods, much less numerous, focus on handling these distribution shifts, for instance by recasting the multi-source learning question as a bayesian multi-task problem [28]. To the best of our knowledge, no work has attempted to handle both the domain shifts that exist between input spaces {𝒳^1^, …, 𝒳^*S*^} and the distribution shifts across {𝒫^1^, …, 𝒫^*S*^}.

The second component of our formalization renders explicit the need to *transfer* knowledge to new subjects. The transferability is warranted by the facts that i) the participants are assumed to be drawn from a homogeneous population, and ii) they have participated in the same experiment, and hence the output space 𝒴 is the same for all individuals. *Experiment 2* has demonstrated that even after multi-source transductive feature standardization, the transfer gap to new subjects remains important. While the construction of anatomical or functional templates provides a technical solution to this problem (see previous paragraph), the existence of this additional gap suggests that given a fixed set of training subjects, the transfer can be further tuned adaptively to the target subject, i.e. differently for two distinct test subjects. This idea is seldomly present in the ISPA literature. We can cite the examples of [45] which introduces a neural code converter specifically tuned at transfering information to a single target subject, [30] which uses classifiers trained on single-subject data to test their transfer capability to all other subjects before stacking them in a second level, or [47] which learns personalized models of emotion perception using transfer component analysis. Our formalization therefore emphasizes the possibility to adaptively take into account the characteristics of the target subject to optimize the transfer of information.

The third component of our formalization, the *transductive* nature of ISPA, offers opportunities to address this transfer question. Indeed, having access to all the samples of the target subject at learning time – without their labels – offers numerous opportunities to perform operations that will minimize the domain and distributions shifts between the training subjects and the target subject. However, this has been exploited in very few ISPA methods; examples include standardizing features subject-by-subject (as discussed above and used in *Experiment 1*), learning a graphical model using all the data of the test subject in [41], or performing joint decision on all data points using a similarity space [33]. But none of these methods actually perform *transductive transfer*, which can also be called *domain adaptation*. Nonetheless, several recent exceptions have appeared in the literature, such as when the maximum mean discrepancy is used to minimize the distribution shift with the target subject [47] or when transductive adaptation is obtained using generative adversarial networks (GANs) [26]. Given the few methods that exploit it and the large beneficial effect observed in *Experiment 1*, making the most of transduction might be a lead to obtain models that generalize better to unseen subjects and in more diverse contexts. The interpretation of such models to perform group inference should however be handled carefully, for instance by checking their consistency across the different folds of the leave-subjects-out cross-validation.

Furthermore, we believe that the three components of our multi-source transductive transfer formalization have never been combined into a single ISPA method. The present paper should therefore be directly useful to pave the road for future methodological work, either by combining existing methods that address the different challenges isolated in this discussion, or by designing new ones. For instance, embedding hyperalignment [18], which has proved efficient to perform multisource representation of multi-subject data, within a probabilistic transductive transfer scheme, such as described in [28], might be an effective way of benefiting from the formalization introduced here. The main challenge with task-based functional neuroimaging data remains the small number of examples available in each subject, which could make most recent machine learning techniques difficult to tune. For this, a parallel could be built between ISPA and another learning problem encountered in neuroimaging and more generally in medicial imaging: multi-site learning, which occurs when accumulating data from patients treated in different hospitals to increase the overall sample size (see initiatives as e.g ENIGMA^1^ or EU-AIMS ^2^). The development of such multi-site studies is of great importance to improve the reliability of CAD systems, but also for questions such as population stratification or lesion segmentation. While in ISPA, the challenge is to overcome inter-individual variability, it is here to handle between-site differences. Model evaluation requires using leavesite-out cross-validation to avoir over-optimistic estimations of the generalization performances [15]. As ISPA, multi-site learning can clearly be formalized within a multi-source transfer setting, but several differences exist: i) in multi-site CAD, one need prediction on single patients, which restrict the choice of learning methods to inductive ones, whereas ISPA can benefit from transduction since it is the prediction of *all* labels corresponding to each of the brain activation maps of a test subject which is informative about neural coding principles; ii) because new patients are continuously being recruited, multi-site learning should favor online learning methods, while ISPA is a purely offline question; iii) labeled samples (i.e patients with their diagnostic) might be available in the target site, which allows exploiting semi-supervized models for multi-site learning while it is not possible for ISPA. Regardless of these differences, the largely overlapping character of the challenges to be met opens numerous opportunities for a convergence of future methodological developments for multi-site learning and ISPA. Lastly, the foreseeable popularization of multi-center task-based functional neuroimaging studies (such as the LEAP project^3^), which should help overcoming the lack of robustness of the results obtained in fMRI, EEG or MEG cognitive neuroscience studies [7], will necessitate to embed our ISPA formalization within a multi-site learning framework, therefore adding an extra level of hierarchy in the framework presented here [15].

## 5. Conclusion

In this paper, we have introduced the first formalization of inter-subject pattern analysis (ISPA) as a multisource transductive transfer learning problem. We have demonstrated the added value of each of the three components of this formalization using proof-of-concept experiments where simple data transformation and learning algorithms were used. Furthermore, we have discussed several opportunities that should be helpful to design new ISPA methods. We hope that this work will contribute to promote ISPA as a multivariate group analysis scheme for task-based functional neuroimaging experiment.

## Acknowledgement

This work was carried out within the *Institut Convergence ILCB* (ANR-16-CONV-0002) and was supported by a grant from the French *Agence Nationale de la Recherche* (ANR-15-CE23-0026).

## Appendix A. Standardization strategies used in Experiment 1

In this section, we provide a thorough description of the different standardization strategies benchmarked in *Experiment 1*. For this, we first define three functions that will be used in each algorithm:

### Algorithm 1 LearnStandardizationParameters

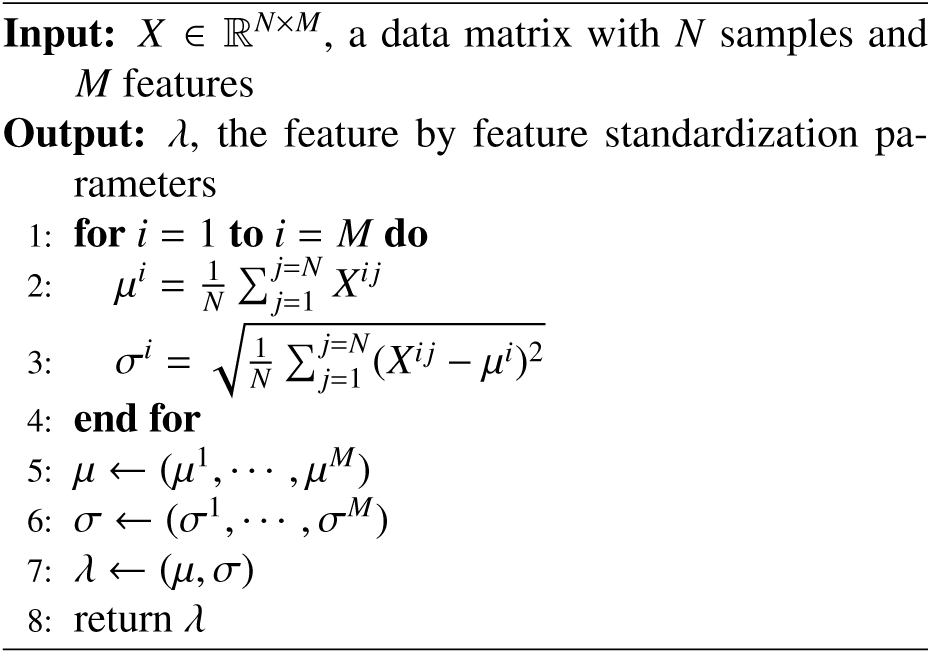

### Algorithm 2 ApplyStandardization

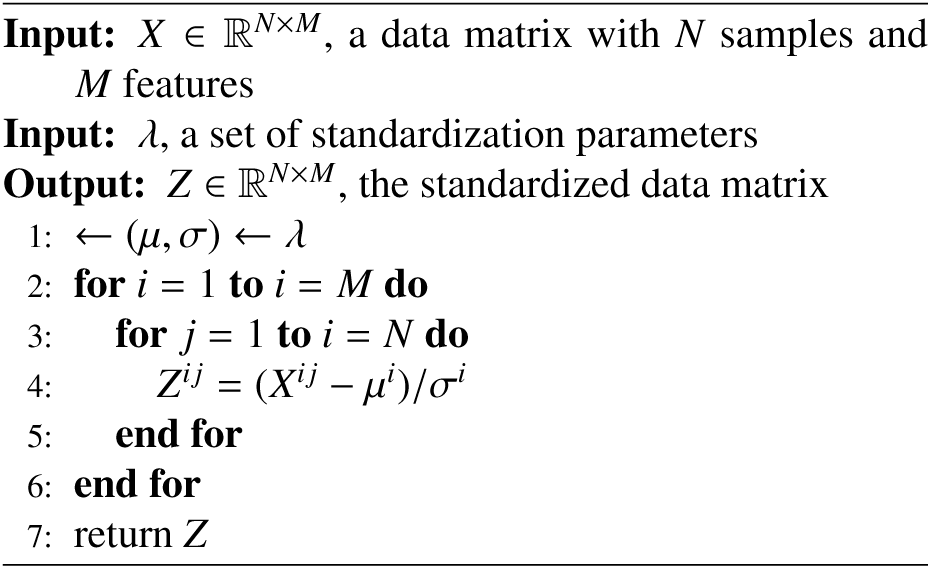

### Algorithm 3 PoolData

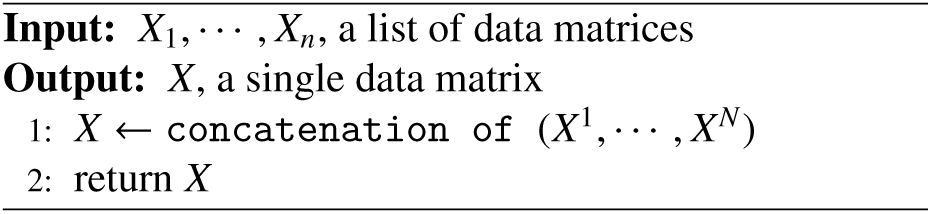

### Appendix A.1. Pooled inductive standardization used in classical machine learning

In classical machine learning, the standardization parameters are estimated on the training set, and used to transform both the training and the test set.

#### Algorithm 4 Pooled inductive standardization

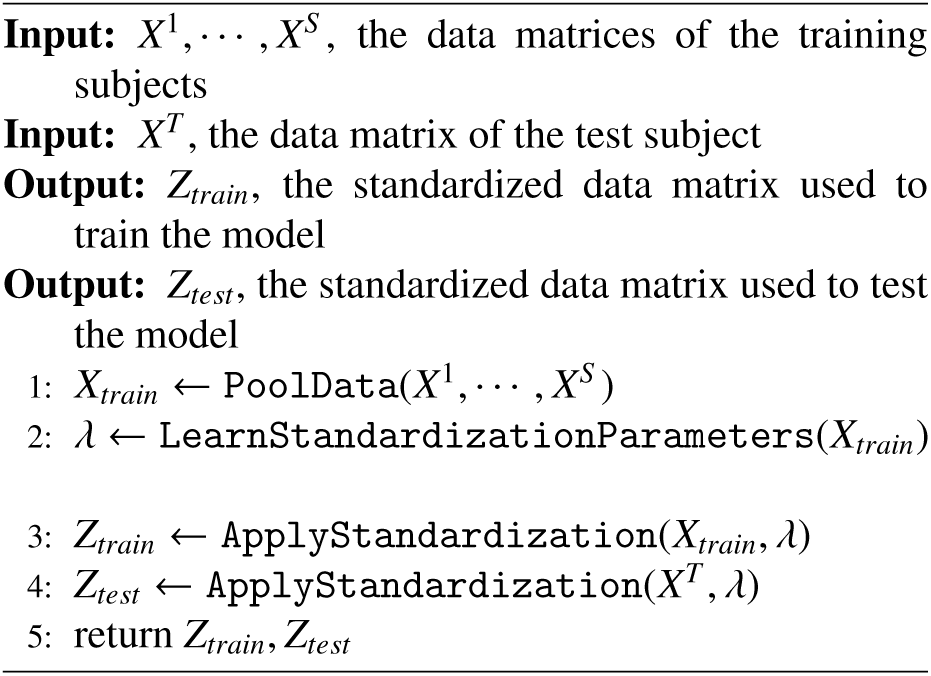

### Appendix A.2. Multi-source transductive standardization

This is the subject-by-subject standardization strategy often encountered in ISPA studies, sometimes without being reported.

#### Algorithm 5 Multi-source transductive standardization

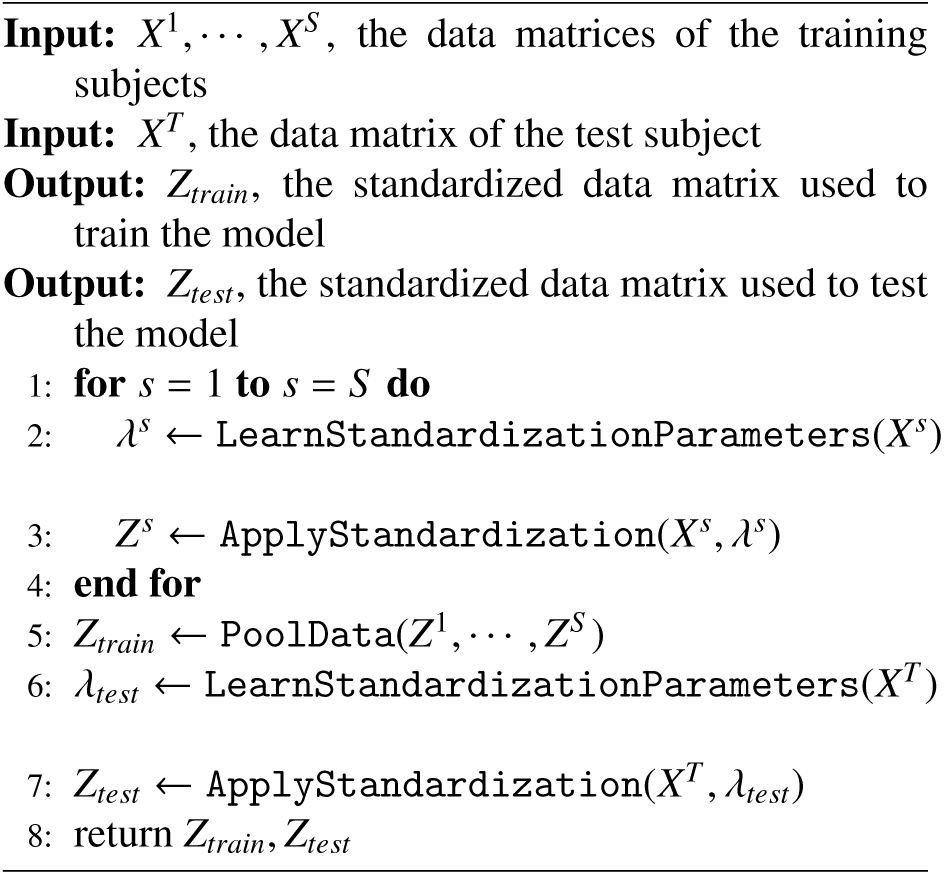

### Appendix A.3. Pooled transductive standardization

This is not a strategy that would be used naturally. We have designed it to allow for distinguishing the effects of the transductive and multi-source settings, thanks to the 2 × 2 factorial design described in *Experiment 1*.

#### Algorithm 6 Pooled transductive standardization

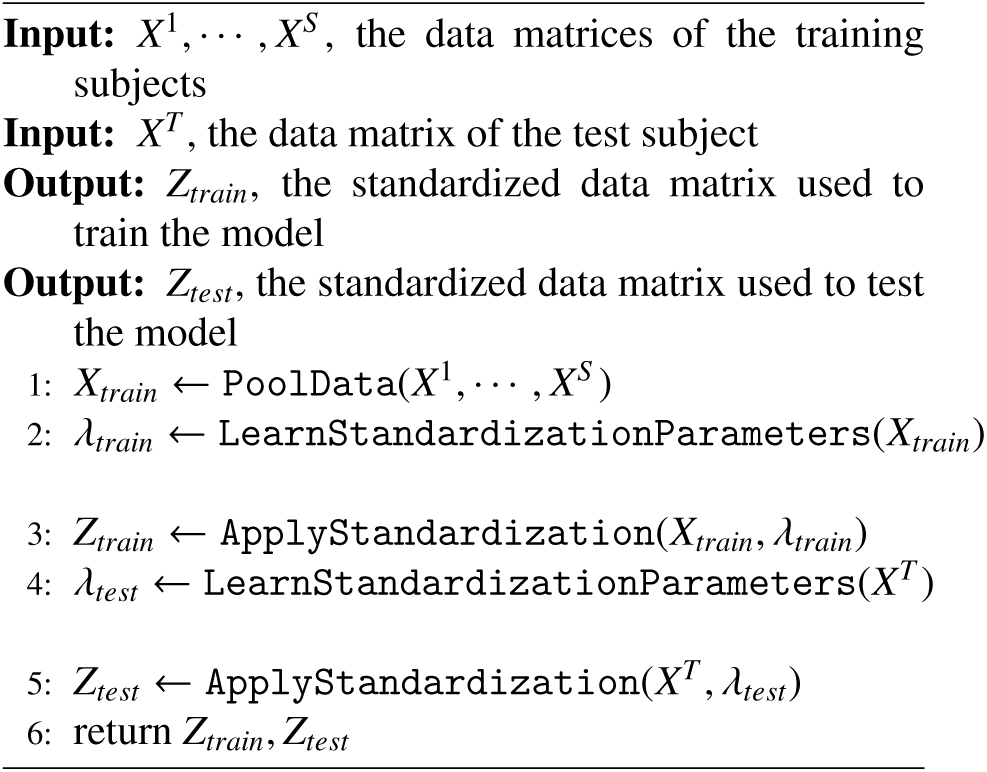

### Appendix A.4. Multi-source inductive standardization

As with the previous strategy, this one would not be used naturally. We have designed it for the same reason, i.e to allow to fullfil the 2 × 2 factorial design described in *Experiment 1*. A challenge here was to select of subject amongst the training ones, without using the test subject. We therefore chose to compute the median training subject, according to its standardization parameters; this is detailed below in a separate algorithm.

#### Algorithm 7 Multi-source inductive standardization

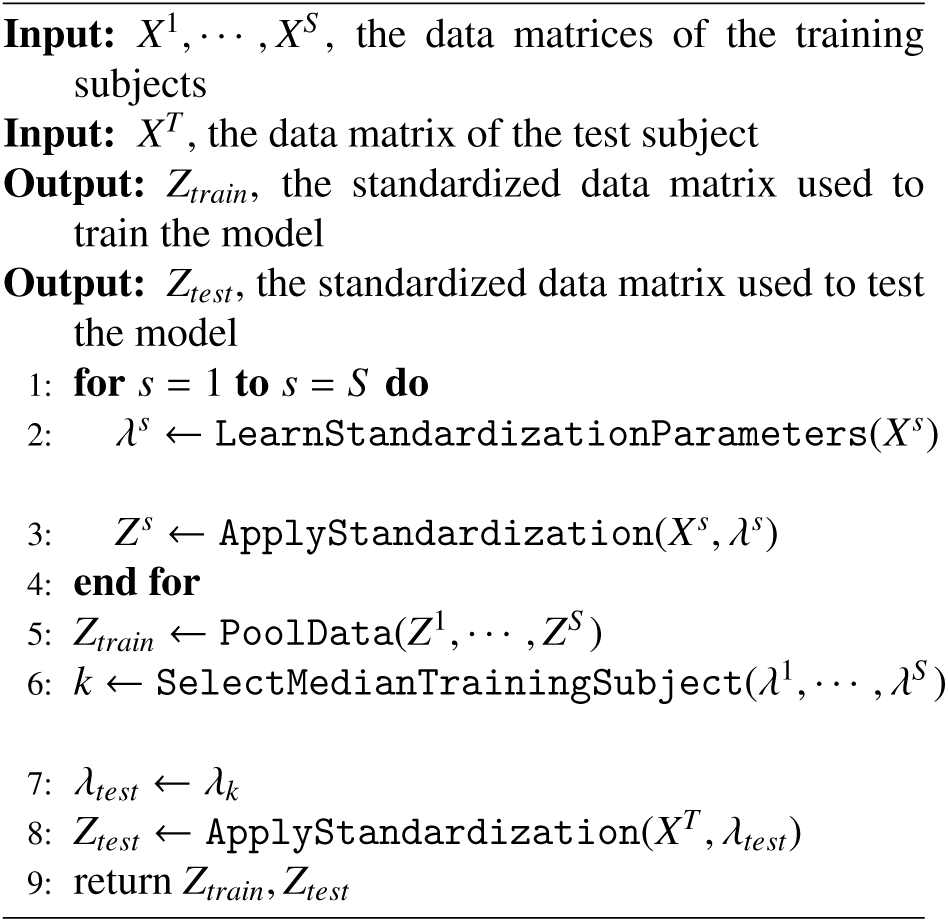

#### Algorithm 8 Definition of the SelectMedianTrainingSubject function used in Algorithm 7

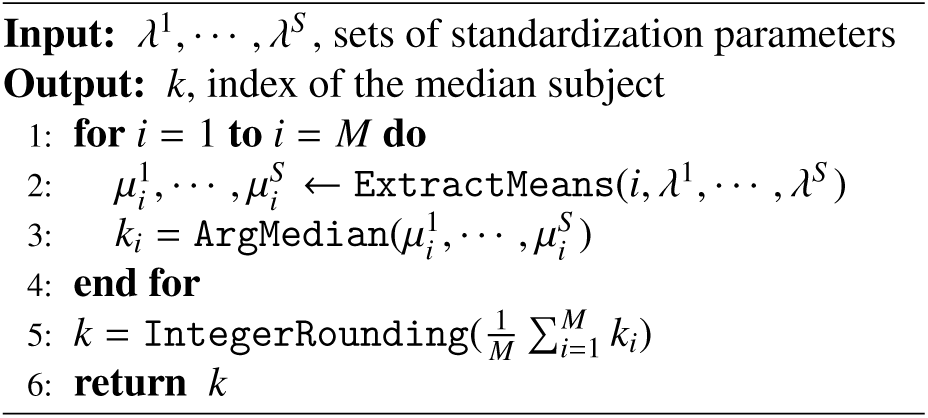

## Footnotes

1 http://enigma.ini.usc.edu/

2 https://www.eu-aims.eu/

3 https://www.eu-aims.eu/the-leap-study

## Notes

### Competing Interest Statement

The authors have declared no competing interest.

